# RNA virus infection reshapes carbon and nitrogen partitioning in a marine diatom

**DOI:** 10.64898/2026.08.30.748135

**Authors:** Emmanuelle Jaouen, Chiara Fiorile, Pascal Riera, Lucie Blondel, Martin Gachenot, Florence Le Gall, Pauline Nogaret, Cédric Leroux, Christophe Six, Sophie Le Panse, Ian Probert, Priscilla Gourvil, Estelle Bigeard, Nathalie Simon, Anne-Claire Baudoux

**Author notes:** Contributed equally to this work (Emmanuelle Jaouen, Chiara Fiorile).

## Abstract

Viral infection is a major yet poorly quantified driver of microbial interactions and biogeochemical fluxes in the ocean. In diatoms, which are key contributors to marine primary production, the extent to which viruses reprogram host cell metabolism and alter elemental cycling remains largely unresolved. Here, we investigated how infection by a lytic single-stranded RNA virus reshapes carbon (C) and nitrogen (N) fluxes in the ecologically relevant nanoplanktonic diatom *Mediolabrus comicus*. Using controlled infection experiments coupled with flow cytometry, electron microscopy, PAM fluorimetry, and stable isotope probing, we resolved infection-driven changes from the population to the cellular scale. Infection induced rapid optical shifts and cellular reorganization, including the formation of membrane-bound viral replication compartments. These changes coincided with early impairment of plastidial functions, as shown by disruption of photosystem II functionality and a concomitant decline in photosynthetic carbon fixation. In contrast, nitrogen uptake was maintained and strongly enhanced during late stages of infection, indicating sustained resource acquisition to support viral replication. This decoupling led to dynamic changes in cellular stoichiometry and, overall, to substantial reductions in population-level carbon and nitrogen assimilation due to growth inhibition. Together, these findings demonstrate that diatom RNA virus infection reshapes host carbon and nitrogen metabolism, with cascading effects on elemental cycling. Our results identify diatom RNA viruses as important drivers of marine biogeochemical processes, with implications for primary production and the fate of organic matter in the ocean.

## Introduction

Diatoms are among the most globally distributed, abundant and diverse photosynthetic protists in the ocean (1,2). These unicellular organisms are central players in ocean biogeochemistry, contributing up to 40% of marine primary production (3–5). Their silica-based frustule makes them dominant contributors to biosilicification, while also providing ballast that drives significant carbon export to the deep ocean (6). Diatoms also contribute substantially to nitrogen cycling, owing to efficient mechanisms for nitrogen uptake and utilization (7,8). Given their major role in biogeochemical cycles, food webs, and carbon export, identifying the processes that regulate diatom-driven fluxes is a central challenge for understanding marine ecosystem functioning (4).

Viruses are increasingly recognized as important mortality agents of marine diatoms (9). Viruses infecting widespread diatom genera have been isolated from different marine environments and typically possess small, non-enveloped capsids (∼30 nm) enclosing positive-sense single-stranded RNA (ssRNA) or single-stranded DNA (ssDNA) genomes (9,10). Most of them exhibit a lytic lifestyle, leading to host cell lysis within days following infection. Environmental surveys indicate that such viruses are globally distributed and actively infect marine diatoms (11–13), yet their consequences for diatom-mediated biogeochemical fluxes remain poorly constrained.

Few studies have examined the fate of organic material released following viral lysis of diatoms. Cellular debris can aggregate into rapidly sinking particles, enhancing particulate carbon export through the viral shuttle (14), while viral lysis also releases dissolved organic matter (DOM) that fuels microbial recycling *via* the viral shunt (15). The composition and bioavailability of this DOM can depend on virus type, with ssRNA and ssDNA diatom viruses differentially influencing its utilization by heterotrophic prokaryotes (15). Thus, viral lysis can shape the partitioning of diatom-derived carbon between export and recycling pathways.

Viruses can also reshape the biogeochemical fate of their hosts before lysis. During infection, the host cell transitions into a distinct state, the *virocell*, characterized by metabolic reprogramming that prioritizes viral replication (16–19). In large DNA virus systems, auxiliary metabolic genes (AMGs) can redirect central carbon and nutrient pathways, altering cellular stoichiometry and energy allocation (16,18). Whether comparable shifts occur during infection of diatoms by small ssRNA and ssDNA viruses, which generally lack AMGs (20), remains largely unexplored. A recent transcriptomic study reported increased expression of genes involved in nitrogen assimilation, photosynthesis, and energy metabolism during infection of *Chaetoceros tenuissimus* by ssRNA and ssDNA viruses (21), suggesting metabolic reprogramming even in the absence of AMGs. However, such transcriptional changes do not necessarily translate into altered rates of carbon and nitrogen assimilation, leaving their consequences for elemental fluxes and biogeochemical cycling unresolved.

The nanoplanktonic diatom *Mediolabrus comicus* (H. Takano) (synonym *Minidiscus comicus,* (22)), provides an ecologically relevant model to address this gap. As one of the smallest known marine centric diatoms (2–8 µm), *M. comicus* is widespread and often dominant in contrasted coastal systems, forming dense blooms that contribute substantially to carbon export (23–26). Although few studies have addressed the regulation of this cosmopolitan diatom, viruses infecting *M. comicus* were detected throughout bloom development in the English Channel (24), indicating recurrent virus–host interactions. Characterized *Mediolabrus* virus exhibit a lytic lifestyle, they belong to the family *Marnaviridae* (ssRNA virus) and distribute among two genera (*Sogarnavirus* and *Kusarnavirus*) (Nogaret et al., in revison). Their global-scale distribution remains to be investigated but considering the ecological significance of *Mediolabrus*, it is likely that viruses associated to these dominant hosts also represent key members of the viral community.

Here, we used controlled infection experiments with *M. comicus* and its lytic RNA virus RCC7291 to quantify virus-induced changes in central biogeochemical processes mediated by diatoms, including photosynthesis and nitrogen assimilation. By focusing on this specific host– virus pair, we extend the virocell framework to RNA virus–diatom interactions and provide novel insights into how viral infection may modulate primary production and elemental cycling in marine ecosystems.

## Material and methods

### Strains and culture conditions

The diatom species *Mediolabrus comicus* RCC4660 was obtained from the Roscoff Culture Collection (RCC, http://www.roscoff-culture-collection.org/). This strain was isolated in 2015 at the SOMLIT-Astan station, in the Western English Channel (48°46’18’’ N, 3°58’6’’ W, (24)) and maintained in K+Si medium (27). Cultures were grown at 18°C, under a 12:12 h light:dark cycle of 100 μmol photons m^−2^ s^−1^ provided by white fluorescent tubes (Philips Master TL_D 18W/865).

The ssRNA virus strain (RCC7291) infecting *M. comicus* was isolated in 2016 from the same sampling site and is also available at the RCC (24). A cryopreserved stock of this viral strain was reactivated by inoculation (1:10 volume) in fresh host culture. The mixture was incubated under host growth conditions until complete host lysis (10 to 15 days). The lysate was collected and filtered over 0.2 µm polyethersulfone syringe filters to be stored at 4°C in darkness.

### Experimental setup

A series of infection experiments was conducted under controlled conditions to investigate the effect of viral infection on host physiology and carbon and nitrogen fluxes. Non-isotope-enriched cultures were used to monitor infection dynamics and host physiology, while parallel isotope-enriched cultures quantified carbon and nitrogen incorporation. This paired design also assessed whether isotope enrichment affected host physiology or infection dynamics (see Results).

A culture of *M. comicus* RCC4660 was grown to 2×10^5^ cells mL⁻¹ and then divided into twelve subcultures of 335 each. Four experimental conditions were established in triplicate: (1) uninfected, non-isotope-enriched; (2) infected, non-isotope-enriched; (3) uninfected, isotope-enriched; and (4) infected, isotope-enriched. Non-enriched media were supplemented with 2 mg L⁻¹ NaH¹²CO₃ and 1 mg L⁻¹ ¹⁴NH₄Cl, whereas isotope-enriched media contained 2 mg L⁻¹ NaH¹³CO₃ (99% ^13^C, Eurisotop) and 1 mg L⁻¹ ¹⁵NH₄Cl (98% ^15^N, Eurisotop). Following substrate amendment, infected cultures were inoculated with 20% (v/v) of virus RCC7291 lysate, which corresponded to a multiplicity of infection of 300, while control cultures received an equivalent volume of culture medium.

All cultures were sampled three times daily over 94 h (at the onset of the light period, 8 h after lights-on, and 15 h after lights-on) to quantify diatom, bacterial, and viral abundances, assess diatom photophysiology, and measure carbon and nitrogen incorporation. Samples for microscopic examination of diatom morphology were collected every 24 h from the non-isotope-enriched treatments only. The analytical procedures are described below.

### Flow cytometry

All flow cytometry analyses were performed at the RECYF platform (https://www.sb-roscoff.fr/fr/plateforme-de-cytometrie-recyf) using a NovoCyte Advanteon flow cytometer (Agilent, Santa Clara, CA, USA) equipped with a 488 nm blue laser.

For diatom population analyses, fresh samples were collected and immediately analyzed for 1 min at a flow rate of 80 μL min⁻¹ setting the trigger threshold on red fluorescence (RFL; >630 nm). Diatom populations were discriminated based on forward scatter (FSC) and RFL signals. These analyses were used to monitor cell counts and assess cell optical properties. For this purpose, the peak heights of FSC, side scatter (SSC), and RFL signals were normalized to internal standard beads (0.95 µm YG beads; Polysciences, Warrington, PA, USA).

For bacterial abundance, analyses were performed on glutaraldehyde-fixed samples (final concentration 0.5%) stored at −80°C until analysis. After thawing, samples were diluted in 0.2 μm-filtered, autoclaved TE buffer (pH 8) and stained with SYBR Green I (final dilution 1:10,000 from the commercial stock) for 15 min at room temperature in darkness. Samples were analyzed for 1 min at a flow rate of 80 μL min⁻¹ setting the trigger threshold on green fluorescence (GFL). Flow cytometry data were processed using NovoExpress software.

### Viral titer

Virus samples were serially diluted (from 10^−1^ to 10^−14^) in exponentially growing host in 48-well plates (1mL, final volume). Triplicate dilution series were incubated under host growth conditions for 3 weeks. Culture clearing was assessed visually, with wells showing lysis scored as positive. Viral titers (infectious viruses mL⁻¹) were then estimated using the most probable number (MPN, (22)) method (US. Environmental Protection Agency; Cornish & Fisher Limits, https://mostprobablenumbercalculator.epa.gov/mpnForm). The latent period and the burst size, were calculated from the dynamics of infectious viruses. The latent period was calculated as the lapse-time between virus inoculation in host culture and the release of infectious viruses in the medium. The burst size (BS) representing the number of virions produced by a lysed cell was calculated as follows:

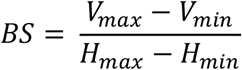

where V and H are virus and host abundances, respectively.

### Pulse amplitude modulated fluorimetry

To assess the impact of viral infection on host photophysiology, the maximal quantum yield of photosystem II (F_V_/F_M_) was measured using a pulse-amplitude-modulated fluorometer (Phyto-PAM II, Walz, Germany). Fresh samples (2 mL) were dark-acclimated for 5 min before measuring basal fluorescence (F_0_) under non-actinic modulated light (440 nm). Maximum fluorescence (F_M_) was then determined after a saturating white-light pulse (5000 μmol photons m⁻² s⁻¹, 400 ms), and F_V_/F_M_ was calculated as:

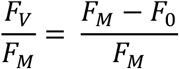

where F_V_ is the variable fluorescence.

To assess the effect of viral infection on light absorption, the photosystem II absorption cross-section, σ(II), was measured. Culture aliquots were exposed to weak far-red light for 2 min before a single-turnover blue-light pulse (440 nm, 2600 μmol photons m⁻² s⁻¹, 80 μs) was applied to record O-I₁ fluorescence kinetics. After verifying saturation, induction curves were modeled and σ(II) for blue light was calculated using PhytoWin-3 software (29).

### Isotopic mass spectrometry

Uninfected and infected culture aliquots (10 mL) were filtered onto pre-combusted (500°C for 4 h) GF/C filters (Whatman) to collect diatom while excluding most viral and bacterial contaminants. Triplicate blanks were prepared using 10 mL of enriched and non-enriched medium. Filters were dried for 48 h at 60°C, folded into tin capsule (12 × 4mm) and analysed at the METABOMER platform (https://www.sb-roscoff.fr/fr/plateforme-de-metabolomique-metabomer).

Cellular ^13^C and ^15^N isotopic composition and content were determined by an elemental analyser (EA Isolink CN/OH, Thermo Fisher) coupled to an isotope ratio mass spectrometer (IRMS, Delta V Advantage, Thermo Fisher) via a Conflow IV open-split interface. Data from the EA-IRMS were processed using ISODAT 3.0 software, Microsoft Excel, and Python (version 3.12). Isotopic values were blank-corrected using mean triplicate blanks and expressed in delta notation (δ, ‰) relative to international reference standards (30,31):

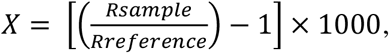

where X is either δ^13^C (carbon) or δ^15^N (nitrogen), R_sample_ is the isotopic ratio ^13^C/^12^C for carbon and ^15^N/^14^N for nitrogen measured in the sample, and R_reference_ is the corresponding isotopic ratio of the international standard (Vienna Pee Dee Belemnite, V-PDB, for carbon and atmospheric N_2_ for nitrogen). Analytical precision, estimated from repeated measurements of laboratory standards (n = 40) was ±0.10‰ for δ^13^C and ±0.15‰ for δ^15^N (n = 40).

To assess isotope incorporation, normalized and blank-corrected ^13^C and ^15^N were converted to isotopic ratios (R) and fractional abundances (F) following (32,33):

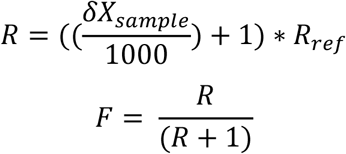

where R is the ratio of either ^13^C/^12^C or ^15^N/^14^N; δX is the normalised-blank-corrected-δ-value of the sample, either δ^13^C or δ^15^N; R_ref_ is the isotopic ratio of the reference standard (R_vpdp_ = 0.0112372 for carbon and R_air_ = 0.0036765 for nitrogen), and F is the fractional abundance of the heavy isotope ^13^C or ^15^N.

Population-level uptake (I_pop_) of the heavy isotope was calculated as:

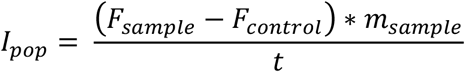

where F_sample_ is the fractional abundance measured in the isotopically enriched treatment, F_control_ is the fractional abundance measured in the corresponding non-enriched control, m_sample_ is the total carbon or nitrogen content of the sample (µg C or µg N), and (t) is the incubation time (h). Uptake rates are expressed as μg ^13^C h^−1^ or μg ^15^N h^−1^.

Cell-specific uptake (Icell) of the heavy isotope was calculated by normalizing population-level uptake to the number of cells determined by flow cytometry:

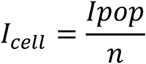

or equivalently,

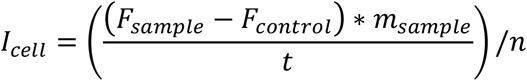

where (n) is the number of cells in the sample. Cell-specific uptake is expressed as µg C cell^−1^ h^−1^ or µg N cell^−1^ h^−1^

The effect of viral infection on isotope incorporation was expressed as the percentage change relative to the uninfected control:

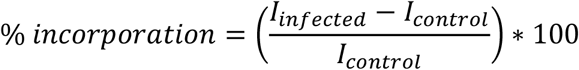

where I_infected_ and I_control_ correspond to the uptake rates either I_pop_ or I_cell_ measured in infected and uninfected cultures, respectively. Positive values indicate enhanced isotope incorporation in infected cultures, whereas negative values indicate reduced incorporation relative to controls.

### Optical and Transmission Electron microscopy

For optical microscopy, a drop of fresh sample was placed between a slide and a coverslip. Cells were then observed under a Nikon 80I microscope using a 100X objective and Nomarski differential interference contrast. Images were captured using an Infinity 3-6UR camera (Teledyne Vision Solutions).

For transmission electron microscopy, samples were fixed in 1% glutaraldehyde (EM-Grade) and 0.01% pluronic acid (F-68, Sigma-Aldrich) for 5 min at 18°C, then centrifuged at 4000 x *g* for 30 min and resuspended in a solution containing 0.4 M Cacodylate pH 7.5, 3.1% glutaraldehyde, 1.8% NaCl in H₂O. After storage at 4°C, samples were post-fixed with 1% OsO₄, (1 h at 4°C), washed in 0.4 M cacodylate, 2% NaCl, dehydrated in an ethanol series (30–100%), embedded in Spurr epoxy resin (Electron Microscopy Sciences) and polymerized for 2 days at 60°C. Thin sections (40 to 70 nm) were obtained with a Leica Ultracut microtome, mounted on copper grids. Sections were contrasted with saturated uranyl acetate and lead citrate according to the Reynolds method and were imaged using JEOL-JEM 1400 TEM (80 kV) at the Merimage platform (https://www.sb-roscoff.fr/fr/plateforme-d-imagerie-merimage).

### Statistical analyses

All analyses were performed using Python 3.12 (Python Software Foundation, 2023).

For cytometric parameters (cell counts, RFL, SSC, FSC), a two-way ANOVA was used to test the effects of infection, enrichment, and their interaction, independently at each sampling time. Residuals from each ANOVA model were tested for normality (Shapiro–Wilk test) and homogeneity of variances (Levene’s test). Only parameters meeting these assumptions (p>0.05) were interpreted directly from ANOVA; otherwise, results were interpreted with caution.

For EA-IRMS data, comparisons between two groups (e.g., enriched infected *vs.* enriched control) were performed using parametric t-tests when data were normally distributed with equal variances. When data failed the normality or equal variance assumptions, non-parametric Mann– Whitney U tests were used. Analyses were repeated across multiple time points to assess temporal changes.

## Results

### Viral infection dynamics of *Mediolabrus comicus*

*M. comicus* grew for 46 h before reaching stationary phase. In infected cultures, growth was inhibited from 13 hours post-infection (hpi), followed by progressive host cell lysis beginning 70 hpi (Fig. 1). A two-way ANOVA confirmed a significant effect of infection on host abundances from 6 hpi (p < 0.05). The viral titer of RCC7291 increased progressively from 46 hpi onward, indicating that infectious virions were released before the onset of host cell lysis. Based on these results, a viral latent period of 46 – 54 h was estimated, and the average burst size was calculated to be 3.6×10⁴ virions cell⁻¹.

**Figure 1.**
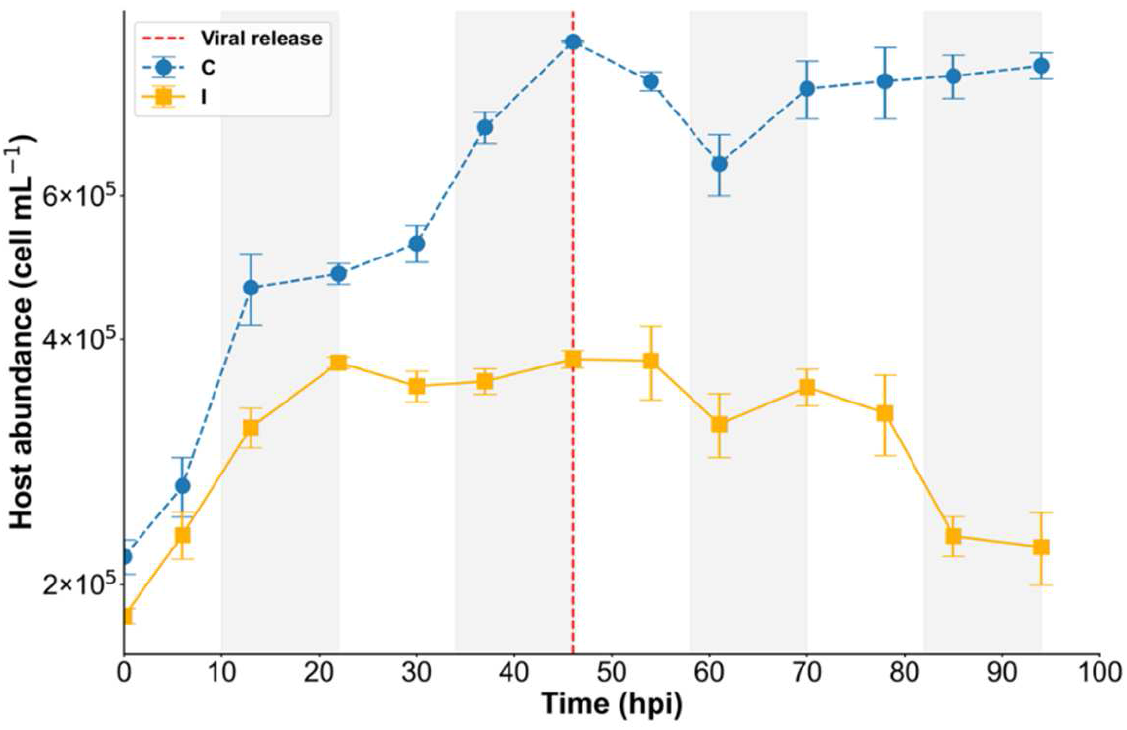
Variation in *Mediolabrus comicus* cell abundance (cells mL⁻¹, logarithmic scale) over the course of the experiment. Control cultures (C) are shown by dashed blue line and Infected (I) cultures by solid yellow line. Values represent mean ± SD (n = 3). Grey boxes indicate night periods. The vertical dashed red line indicates the estimated timing of virus release, determined by plaque assay (Fig. S4b).

### Morphology and optical properties of *Mediolabrus comicus*

Viral infection induced marked morphological and subcellular remodeling in *M. comicus*. Flow cytometry revealed significant alterations in cellular optical properties early during infection. From 37 hpi onward, infected cells displayed increased RFL, proxy for cellular chlorophyll content, and FSC, proxy for cell biovolume, relative to controls, and both parameters then remained elevated throughout the time course (Fig. S1a,b). SSC, reflecting intracellular structural complexity, increased progressively and reached approximately a twofold increase in infected populations by the end of the experiment (Fig. S1c). Optical microscopy observations further indicated an increase in cell roundness within 22 hpi (Fig. S2). Two-way ANOVA confirmed a significant effect of infection on all measured parameters (p < 0.05).

TEM analyses provided further insights into the ultrastructural basis of the cytometric shifts observed during infection (Fig. 2). Control cells exhibited a preserved ultrastructure, including a well-defined nucleus, plastids with organized thylakoid membranes, mitochondria, vacuoles, and pyrenoids (Fig. 2a). Infected cells (virocells) underwent progressive structural alterations. Until 46 hpi, most cells retained their overall integrity but exhibited increased shape heterogeneity, accumulation of vesicular structures, and localized thylakoid disorganization (Fig. 2b,c). By 70 hpi, damaged cells and empty frustules were frequently observed. Cells that retained an overall preserved morphology contained three types of cytoplasmic inclusions: (i) membrane-bound electron-dense compartments containing many viral particles arranged in crystalline arrays and interpreted as viral assembly-like compartments; (ii) numerous vesicle-like structures distributed throughout the cytoplasm; and (iii) an apparent increase in lipid droplets (Fig. 2d,e). In parallel, plastids appeared rearranged but remained present and were sometimes abundant (Fig. 2e). These infection-associated structural features became increasingly pronounced from 70 hpi onward and remained prevalent until the end of the experiment.

**Figure 2.**
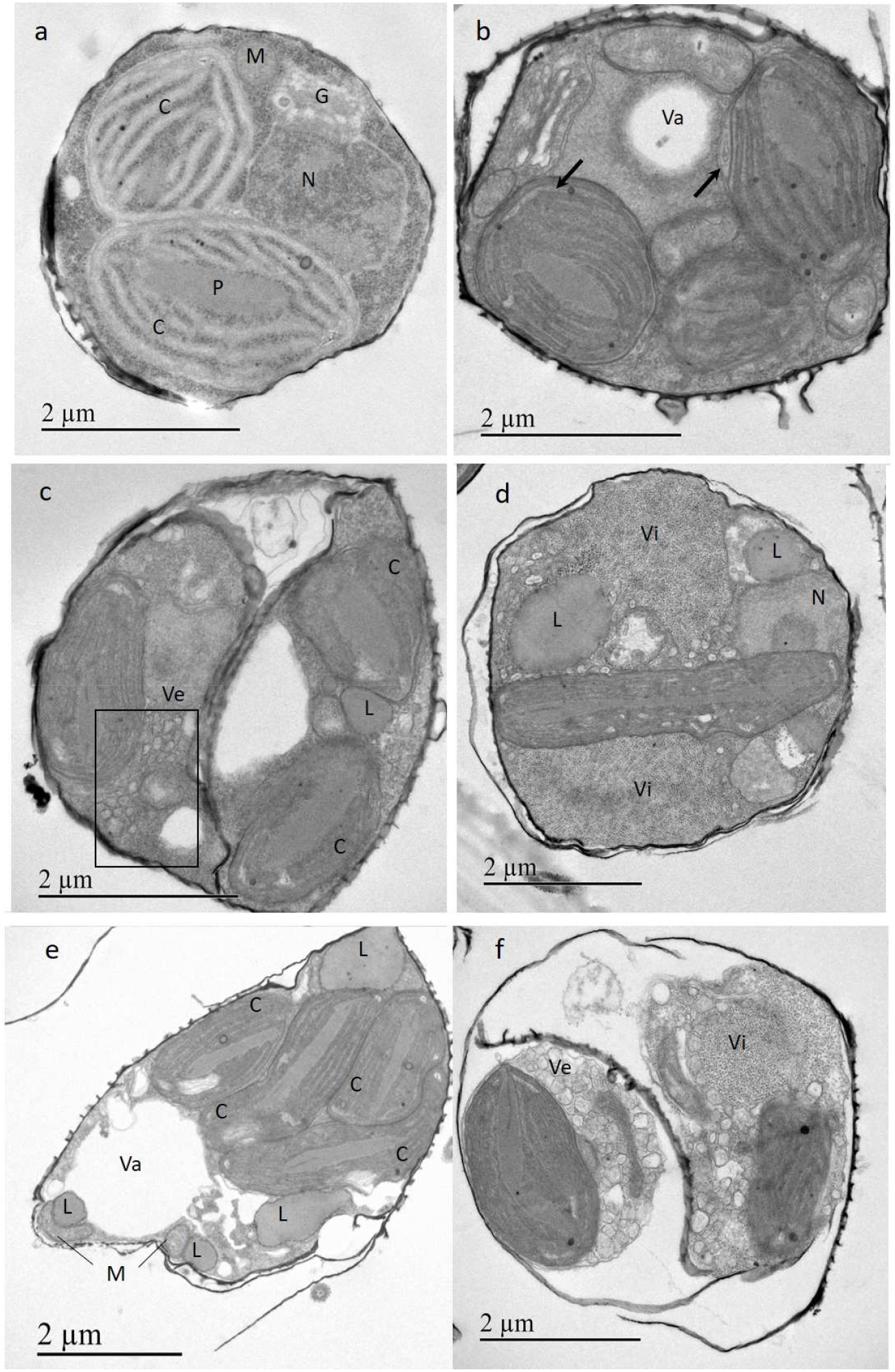
Ultrastructural changes in *Mediolabrus comicus* cells during the course of infection assessed by transmission electron microscopy. **(a)** Control cells showing intact morphology and preserved organelle integrity. **(b–f)** Infected cells displaying progressive alterations. **(b, c)** Cells at 24 and 48 hpi, exhibiting heterogeneous shapes, plastid deformation (arrow), and numerous vesicles. **(d, e)** Cells at 70 hpi, exhibiting viral assembly–like compartments, an increased number of plastids, and lipid droplets. **(f)** Cells at 94 hpi, partially occupying the frustule volume and showing extensive cytoplasmic reorganization, with prominent viral compartments and numerous vesicles. Nucleus (N), chloroplast (C) containing the pyrenoid (P), mitochondrion (M), Golgi apparatus (G), vacuole (Va), vesicle (Ve), viral assembly–like compartment (Vi), and lipid droplet (L).

### Photosystem II functionality

Optical and morphological shifts coincided with alterations in photosynthetic performance. The maximal quantum yield of photosystem II (F_V_/F_M_), reflecting the functionality of photosystem II, ranged between 0.57 and 0.63 in control cultures, exhibiting consistent day-night oscillations during the experiment course. In infected cells, this circadian pattern was disrupted from 30 hpi, and F_V_/F_M_ declined sharply from 46 hpi (Fig. 3a). By 85 hpi, F_V_/F_M_ in infected cells was reduced by approximately twofold relative to controls, averaging 0.29. The functional absorption cross-section of photosystem II, σ(II), decreased from late exponential phase onward (37 h) in the control cultures. In infected cells, it was altered from 6 hpi in infected cells, indicating that light absorption capacity of photosystem II was affected at early stages of infection, well before viral release and host burst (Fig. 3b).

**Figure 3.**
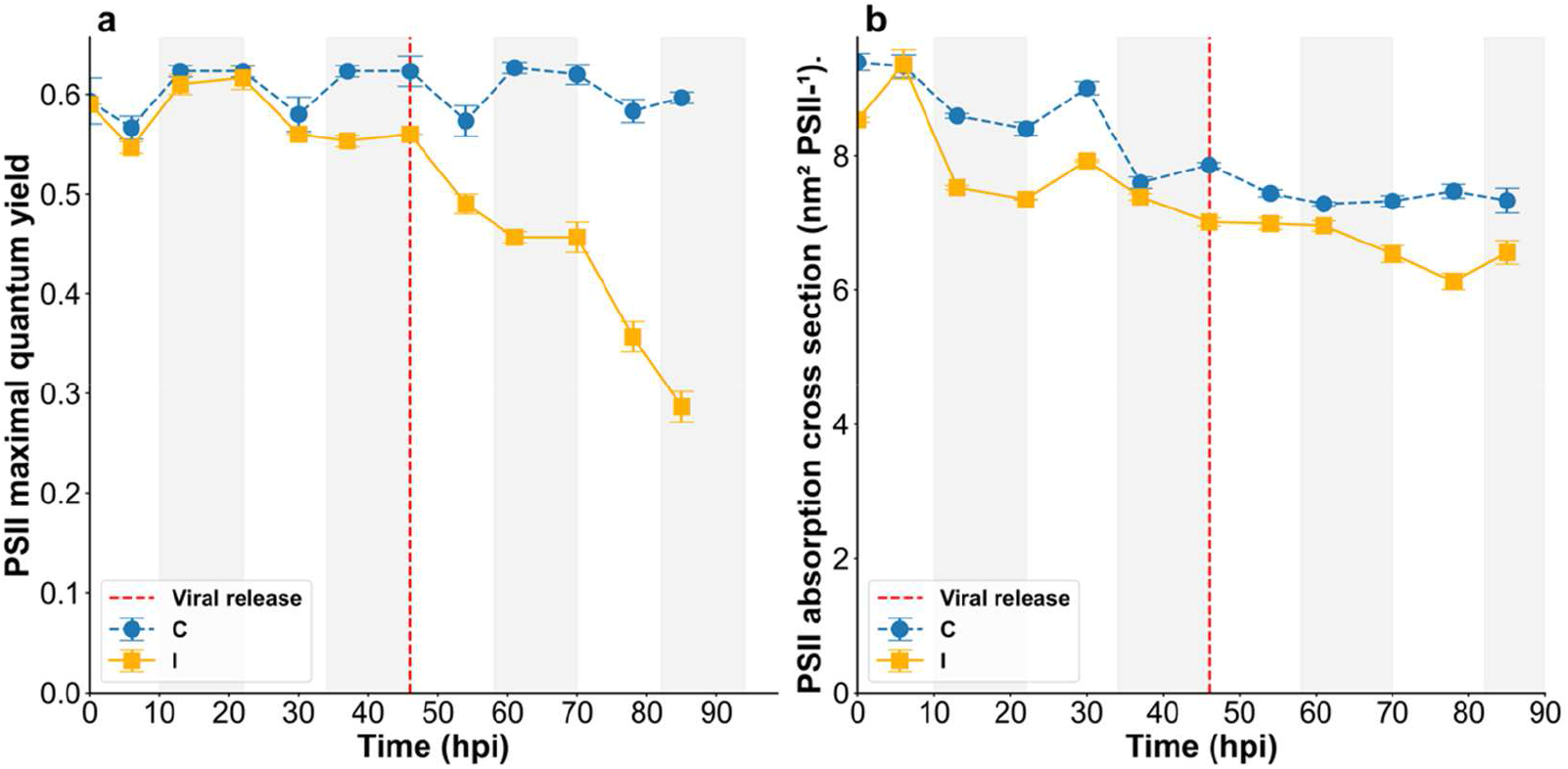
Variation in photosystem II (PSII) parameters of *Mediolabrus comicus* during infection assessed by PAM fluorimetry. **(a)** PSII maximal quantum yield (F_V_/F_M_), proxy of PSII functionality. **(b)** Functional absorption cross-section of PSII (σ(II)) at 440 nm. Control cultures (C) are shown by dashed blue line and Infected cultures (I) by solid yellow line. Values represent mean ± SD (n = 3). Grey boxes indicate night periods. The vertical dashed red line indicates the estimated timing of virus release, determined by plaque assay (Fig. S4b).

### Cellular carbon to nitrogen mass ratio

Initial cellular carbon and nitrogen content amounted to 159 pg C cell^−1^ and 22 pg N cell^−1^ leading to a C:N mass ratio of 5.6 in controls (Fig. S3, Fig. 4). This ratio changed dynamically during infection. In the non-infected control, it remained relatively stable, fluctuating within a narrow range (5.0-6.2; mean 5.8 ± 1.02) throughout the experiment (Fig. 4). In infected cultures, the overall mean C:N ratio (5.6 ± 0.93) was comparable to that of the controls; however, a higher variance (0.76) was observed compared to control cultures (0.12). Until 30 hpi, C:N ratios were consistently higher in infected cultures, reaching a mean peak value of 6.9. This was followed by a sharp decline between 37–46 hpi, with C:N ratios dropping to 4.3, coinciding with a marked increase in cellular nitrogen content in infected cells while carbon content remained similar in both treatments (Fig. S3a). Thereafter, the C:N ratio gradually increased, converging toward control levels by the late stage of infection (78–85 hpi).

**Figure 4.**
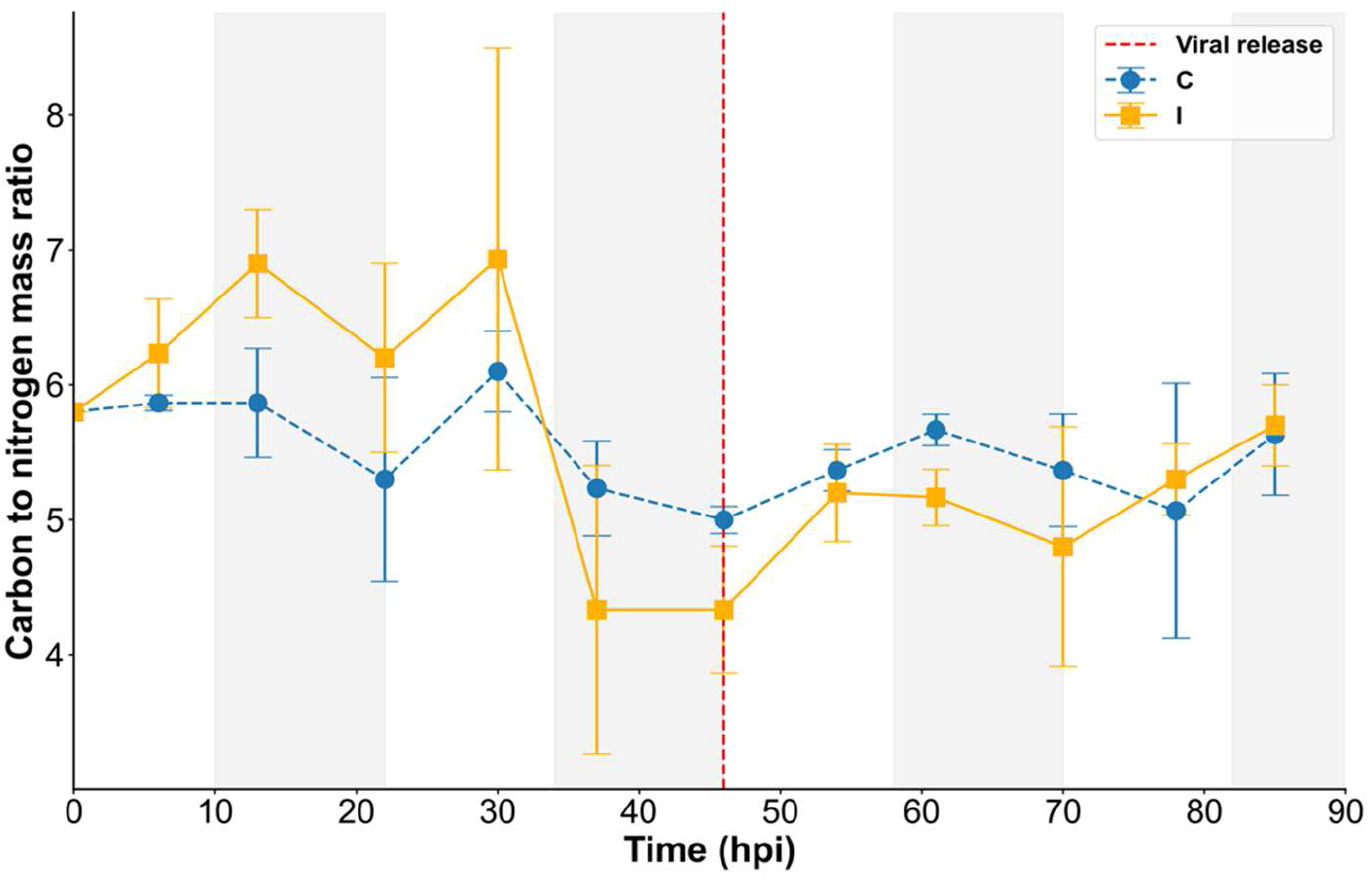
Variation of the carbon to nitrogen mass ratio (C:N) of *Mediolabrus comicus* during infection assessed by mass spectrometry. Control cultures (C) are shown by dashed blue line and Infected cultures (I) by solid yellow line. Values represent mean ± SD (n = 3). Grey boxes indicate night periods. The vertical dashed red line indicates the estimated timing of virus release, determined by plaque assay (Fig. S4b).

### Carbon and nitrogen uptake

To quantify the nutrient uptake rates underlying these stoichiometric shifts, we used ^13^C and ^15^N stable isotope probing to track carbon and nitrogen fluxes during infection, after first confirming that isotopically enriched substrates did not affect host growth, infection dynamics, and optical properties (Fig. S4, Table S1). Overall, enriched treatments were indistinguishable from their corresponding non-enriched controls. Although viral titers appeared to increase more slowly in the enriched treatments (Fig S4b), infected host dynamics were highly similar across treatments, indicating that this apparent delay most likely reflects the inherent variability of the MPN enumeration method rather than a biological effect.

Having established that isotope enrichment did not alter infection dynamics, we next examined isotope incorporation. The marked increase in δ¹³C values observed in both infected and control cultures amended with ¹³C-enriched bicarbonate confirmed the incorporation of this enriched substrate (Fig. 5a). Differences between conditions became apparent after 13 hpi, when infected cultures exhibited significantly lower δ¹³C enrichment than controls (p < 0.01). From 30 hpi onward, δ¹³C values plateaued in both conditions. Nevertheless, the lower incorporation observed in infected cultures persisted throughout the experiment, with δ¹³C values reaching 544.7‰ at 85 hpi compared with 678.9‰ in controls, consistent with reduced ¹³C incorporation by infected hosts (Fig. 5a).

**Figure 5.**
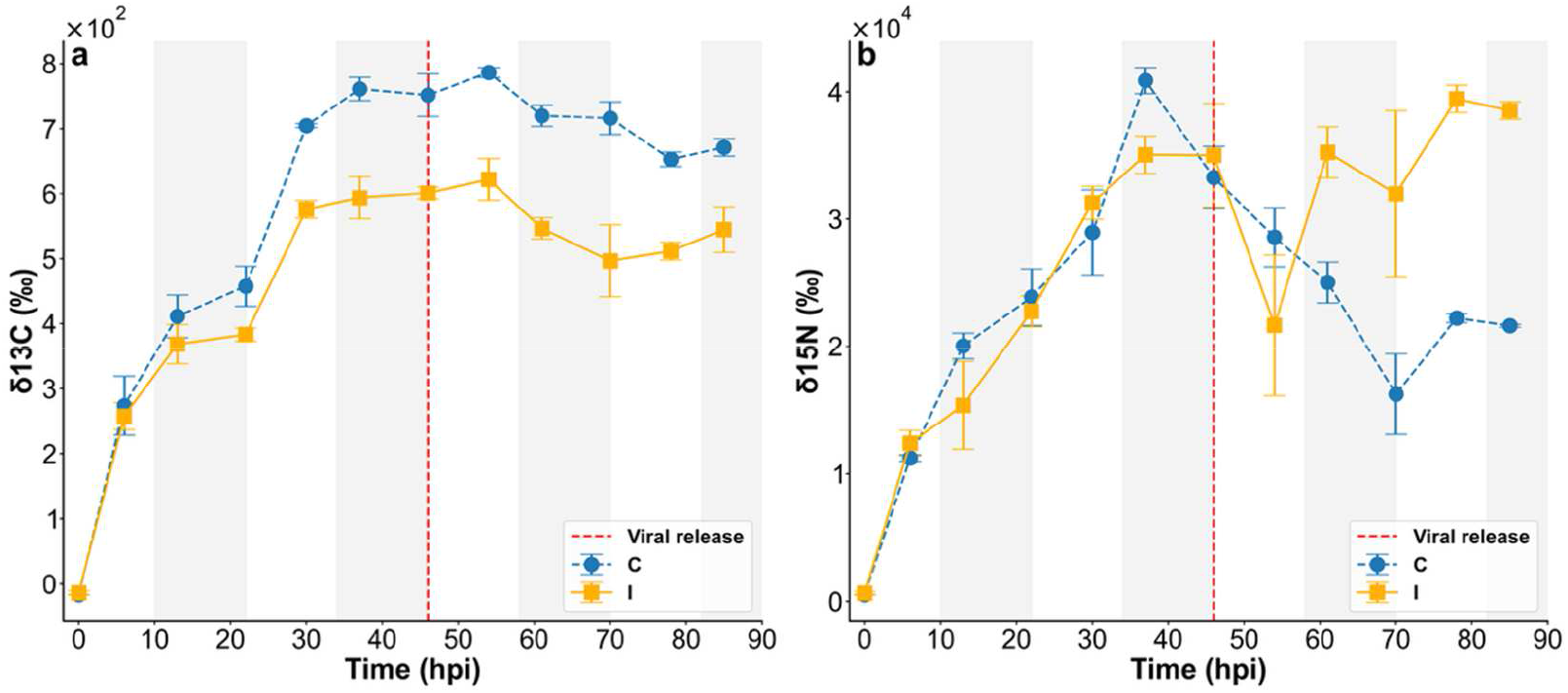
Variation of **(a)** δ¹³C and **(b)** δ15N (expressed as ‰) during infection of *Mediolabrus comicus* amended with ^13^C-enriched NaH¹³CO₃ and ^15^N-enriched ¹⁵NH₄Cl assessed by isotope ratio mass spectrometry. Control cultures (C) are shown by the dashed blue line and Infected cultures (I) by the solid yellow line. Values represent mean ± SD (n = 3). Grey boxes indicate night periods. The vertical dashed red line indicates the estimated timing of virus release, determined by plaque assay (Fig. S4b).

By comparison, δ¹⁵N increased sharply until 37 hpi, reaching approximately 40000‰ in control cultures and 35000‰ in infected cultures (Fig. 5b). This indicates efficient ^15^N incorporation under both conditions despite the reduced growth of infected cultures over this time period (Figs. 1 and 5b). After 37 hpi, the trajectories diverged: δ¹⁵N enrichment declined sharply in controls, whereas in infected cultures it transiently decreased before increasing again. From 61 hpi onward, infected cultures exhibited substantially higher δ¹⁵N enrichment than controls, with a maximum difference of 19426‰ at 70 hpi.

Time-integrated ¹³C and ¹⁵N incorporation rates were calculated from isotopic enrichment data, and infection-induced changes in C and N fluxes were quantified at both the population (bulk; Fig. 6, Fig S5a) and cellular (Fig. 7, Fig. S5b) levels. In control populations, ¹³C fluxes ranged from 0.9×10⁴ to 2.2×10⁴ pg ^13^C h⁻¹ and displayed diel oscillations throughout the experiment (Fig. 6a). Infected populations exhibited substantially lower ¹³C fluxes, ranging from 0.4×10⁴ to 1.3×10⁴ pg ^13^C h⁻¹, and progressively declining throughout the experiment, reaching a maximal reduction of ∼74% relative to controls at the end of the incubation period (Fig. S5a).

**Figure 6.**
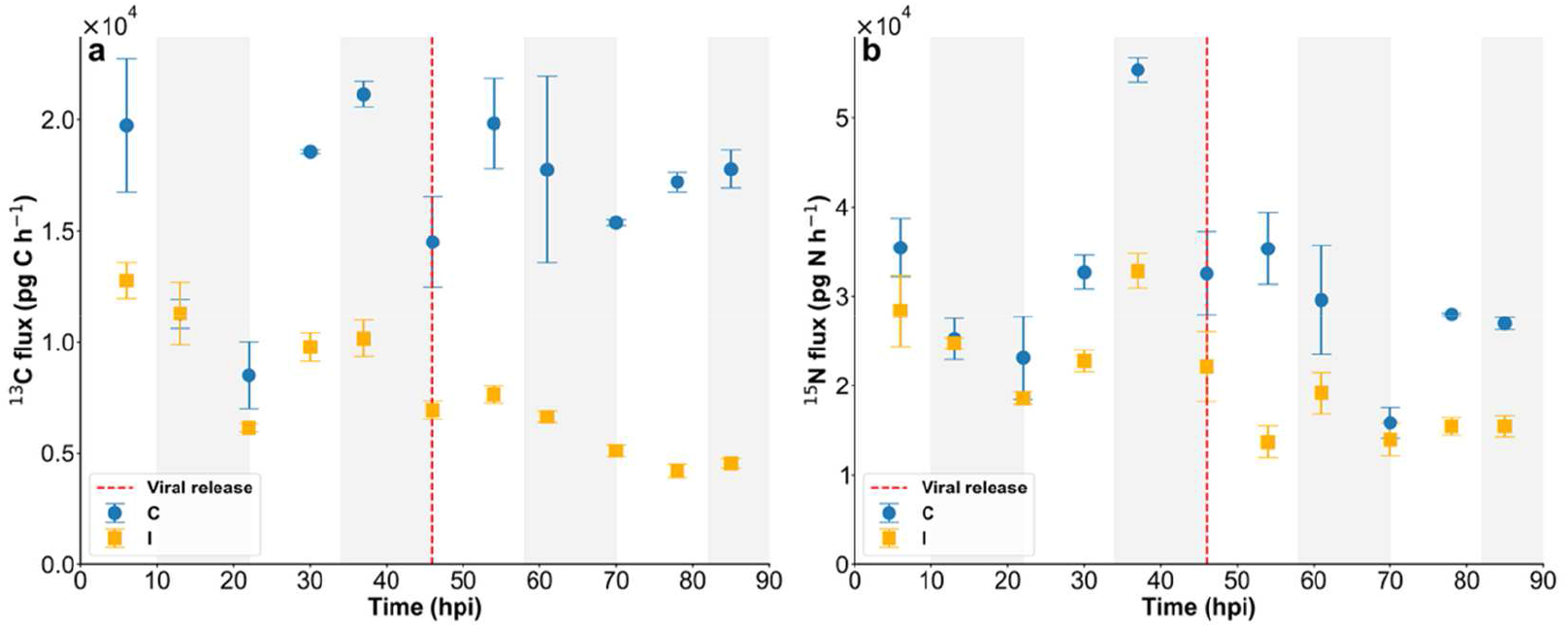
Time-integrated bulk fluxes of **(a)** ¹³C and **(b)** ¹⁵N (pg C or N h⁻¹) during infection of *Mediolabrus comicus* amended with ^13^C-enriched NaH¹³CO₃ and ^15^N-enriched ¹⁵NH₄Cl assessed by isotope ratio mass spectrometry. Control cultures (C) are shown by the dashed blue line and Infected cultures (I) by the solid yellow line. Values represent mean ± SD (n = 3). Grey boxes indicate night periods. The vertical dashed red line indicates the estimated timing of virus release, determined by plaque assay (Fig. S4b).

**Figure 7.**
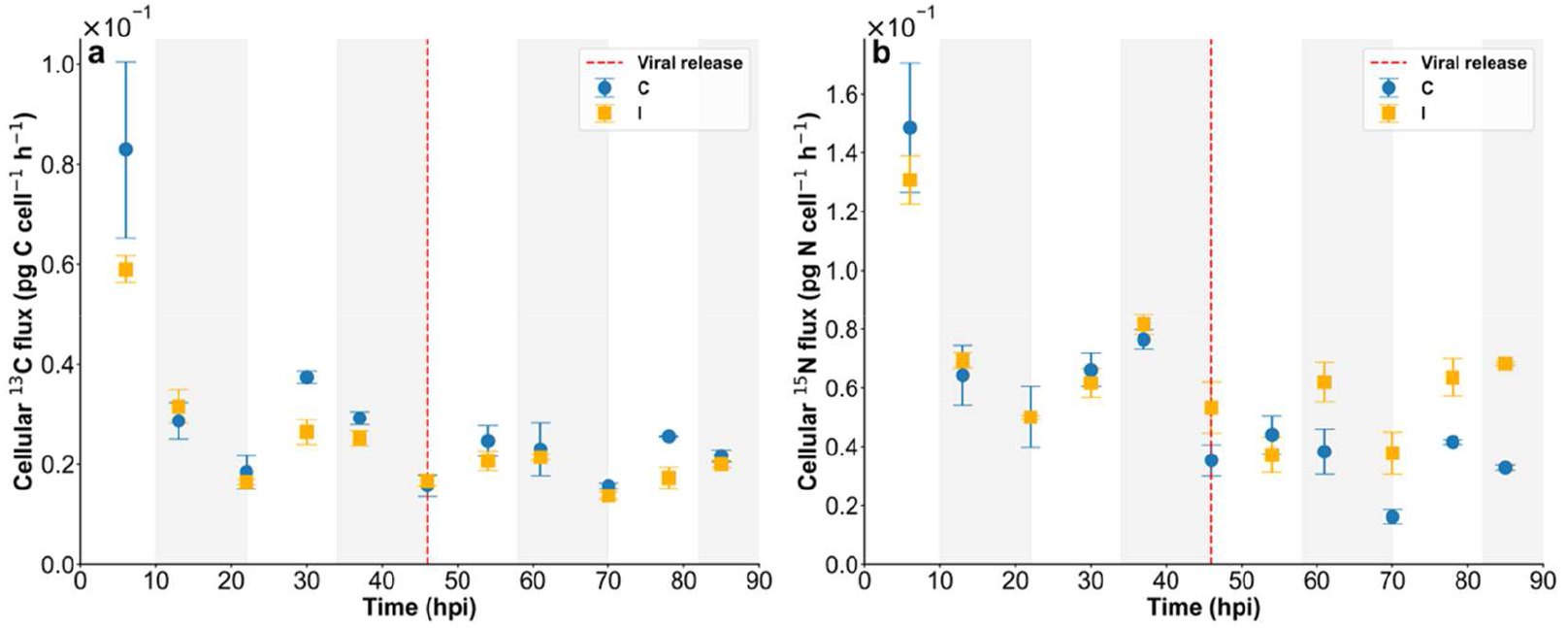
Time-integrated cellular fluxes of **(a)** ¹³C and **(b)** ¹⁵N (pg C or N cell^−1^ h⁻¹) during infection of *Mediolabrus comicus* amended with ^13^C-enriched NaH¹³CO₃ and ^15^N-enriched ¹⁵NH₄Cl assessed by isotope ratio mass spectrometry. Control cultures (C) are shown by the dashed blue line and Infected cultures (I) by the solid yellow line. Values represent mean ± SD (n = 3). Grey boxes indicate night periods. The vertical dashed red line indicates the estimated timing of virus release, determined by plaque assay (Fig. S4b).

Fluxes of ¹⁵N also exhibited dynamic changes, with a peak of 5.7×10⁴ pg ¹⁵N h⁻¹ observed at 37 hpi in control cultures (Fig. 6b). During the early stages of infection, ^15^N fluxes were comparable between infected and control populations. Marked differences, however, emerged from 30 hpi onward, with infected populations displaying consistently lower, yet still dynamic, ¹⁵N fluxes that ultimately reached a ∼44% reduction relative to controls (Fig. S5a). These results indicate that infection profoundly modulates both C and N fluxes at the population scale, resulting in a sustained suppression of C incorporation and a temporally variable, but ultimately reduced, N incorporation.

To gain insight into elemental uptakes at cellular level, bulk fluxes were normalized to diatom abundance (Fig. 7, Fig. S5b). Overall, infected cells incorporated less carbon than control cells (Fig. 7a), with more pronounced decreases in ¹³C fluxes during the light period, reaching maximal reductions of ∼30% at 30 hpi and 78 hpi (Fig. S5b). In contrast, ¹⁵N incorporation showed a distinct temporal pattern. During the early stages of infection (up to 37 hpi), ¹⁵N incorporation rates were comparable between infected and control cells ranging from 0.05-0.13 and 0.05-0.15 pg ^15^N cell⁻¹ h⁻¹. From 46 hpi onward, infected cells exhibited substantially higher ¹⁵N incorporation than controls, reaching a maximum increase of ∼135% at 70 hpi, which, in absolute terms, corresponds to an increase of 0.04 pg ¹⁵N cell⁻¹ h⁻¹ (Fig. 7, and S5b). These cellular scale patterns indicate a decoupling of carbon and nitrogen metabolism during infection, with reduced carbon incorporation—particularly under light conditions—and enhanced nitrogen incorporation at later stages.

## Discussion

Nanoplanktonic diatoms are increasingly recognized as important contributors to marine biogeochemical cycles (34). Although viruses have been shown to interact with these organisms during bloom development in the Western English Channel (24), few viral isolates have been characterized to date, and the cellular and biogeochemical consequences of infection remain poorly understood. Using the ubiquitous nanoplanktonic species *M. comicus* and an associated RNA virus as a model system, we combined ultrastructural, physiological, and elemental analyses to investigate how viral infection alters host cellular organization and metabolic processes relevant to biogeochemical cycling.

The *M. comicus* RNA virus strain RCC7291 is one of the few cultured representatives of the genus *Kusarnavirus* within the family *Marnaviridae* (Nogaret et al., in revision; 35,36). As reported for other diatom-infecting RNA viruses, RCC7291 follows a lytic replication strategy, with viral progeny accumulating in the host cytoplasm and host lysis occurring within 3–4 days post-inoculation (9). Despite the small size of the *M. comicus* host, its burst size was comparable to that reported for viruses infecting larger diatoms, including CtenRNAV type I infecting *C. tenuissimus* and GdelRNAV infecting *Guinardia delicatula* (37,38), highlighting the efficient replication of RCC7291.

Infection rapidly inhibited host growth and induced pronounced morphological and subcellular changes. From 22 hpi onward, transmission electron microscopy revealed numerous vesicle-like structures in the cytoplasm, consistent with viral replication complexes (VRCs). Such structures are characteristic of positive-sense RNA viruses and provide protected microenvironments for viral genome replication, often through the recruitment of membranes from different cellular compartments, including the endoplasmic reticulum, Golgi apparatus, vacuoles, and plastids (39). In *M. comicus*, these vesicle-like structures were frequently observed close to plastids and occasionally associated with localized membrane deformation, suggesting a possible plastidial contribution to their formation. Similar plastid-associated RNA-rich structures have been observed during infection of *Guinardia delicatula* (40). Infection was also associated with increases in FSC and SSC, consistent with cell enlargement, increased internal complexity, and rounding, further indicating extensive remodeling of host cellular organization.

These cellular changes were accompanied by shifts in elemental stoichiometry. Uninfected cells maintained relatively stable C:N mass ratios (5.0–6.1) within the range reported for diatoms (41), despite variation in cellular C and N quotas over the incubation period (148 ± 44.2 pg C cell⁻¹ and 26 ± 8.2 pg N cell⁻¹). Due to the limited availability of cultured nanoplanktonic diatoms, elemental quota data for this size class remain scarce. *M. comicus* cellular quotas of were 10-to 20-fold higher than those reported for a similarly sized *Nitzschia* species (41), potentially reflecting differences in strain, physiology, or methodology. In contrast, virocells exhibited more variable C:N ratios (4.18–7.0), particularly between 30 and 54 hpi, when cellular N quotas increased concomitantly with the probable accumulation of nitrogen-rich viral progeny observed by transmission electron microscopy. These changes suggest that viral replication alters host elemental allocation before cell lysis.

Time-resolved stable isotope measurements further revealed a pronounced divergence between carbon and nitrogen assimilation. Nitrogen incorporation in virocells remained comparable to controls until 37 hpi, indicating sustained N assimilation despite infection-induced growth inhibition. From 46 hpi onward, N incorporation patterns diverged: ¹⁵N enrichment in controls, which had entered stationary phase, declined sharply—suggesting the excretion of nitrogen-rich compounds and/or reduced uptake—, whereas infected cells maintained N incorporation, albeit with increased variability. A pronounced drawdown in infected cultures coincided with increasing viral concentrations in the medium, consistent with the production and release of ¹⁵N-enriched viral particles. Together with previous reports of upregulated N assimilation genes during infection of *C. tenuissimus* diatom (21), the sustained N assimilation throughout infection likely reflects the high N demand associated with viral protein and genome synthesis. As reported previously for DNA viruses (42–44), these results suggest that extracellular N sources contribute substantially to this demand, rather than viral production relying solely on N recycled from host biomass.

In contrast to nitrogen, ¹³C incorporation followed a distinct trajectory, indicating marked decoupling of C and N fluxes at the cellular level during infection. Carbon assimilation was consistently reduced in virocells relative to controls from the early stages of infection, coinciding with a progressive decline in photosynthetic performance. This reduction was accompanied by a progressive decline in the photosystem II antenna size, followed by a decreased reaction center functionality prior to viral release, indicating a reduced capacity for light-driven energy conversion. However, the gradual rather than abrupt decline in photosynthetic activity suggests that photosynthetic processes were impaired but not completely suppressed, as reported for HaRNAV infecting the raphidophyte *Heterosigma akashiwo* (45). Thus, infected cells retain some capacity for carbon assimilation and photosynthetic energy production during infection, potentially contributing carbon and energy to support viral production.

Despite the progressive decline in PSII functionality and carbon assimilation, cellular chlorophyll fluorescence was higher in virocells than in controls throughout the experiment. This increase may reflect higher cellular chlorophyll content, as suggested by the presence of numerous plastids in some virocells. A similar response may occur in other diatom–virus systems, as Hongo and Tomaru (21) reported increased expression of genes involved in chlorophyll biosynthesis during infection of the diatom *C. tenuissimus*. In our system, infection-induced growth arrest may contribute to this pattern by limiting the redistribution of chlorophyll and plastid content associated with cell division. Alternatively, enhanced fluorescence may reflect altered partitioning of absorbed excitation energy, potentially as part of a photoprotective response. These observations suggest that infection does not simply suppress photosynthesis but alters the coupling between light harvesting, photochemical energy conversion, and carbon fixation.

The reduction in carbon assimilation raises the question of how the remaining assimilated carbon is redistributed during infection. Beyond photosynthetic carbon fixation *via* the Calvin cycle, inorganic carbon can be incorporated into other biosynthetic pathways, including *de novo* fatty acid synthesis, which supports membrane biogenesis and storage lipid production (46). Infection could therefore alter the allocation of assimilated carbon among competing cellular sinks, potentially redirecting carbon toward membrane production, required for the formation of viral replication structures, or toward storage compounds, which could explain the apparent accumulation of lipid droplets. Resolving these alternative carbon sinks will require molecular and metabolic approaches capable of tracking carbon allocation beyond bulk photosynthetic fixation.

Taken together, the pronounced decoupling of C and N fluxes at the cellular level, combined with reduced growth, resulted in substantial population-level losses in elemental incorporation. By the end of the experiment, C and N uptake were reduced by 74% and 44%, respectively, in infected cultures. Given the central role of diatoms in marine biogeochemical cycling, infection-induced changes in C and N allocation could alter the quantity and stoichiometry of organic matter available to the microbial loop. Extending the virocell framework to RNA viruses, our results demonstrate that viral infection can substantially reshape diatom elemental metabolism, with potential consequences for ecosystem-level biogeochemical fluxes.

## Concluding remarks

Growth rate, cell size, and elemental stoichiometry are key traits for tracking phytoplankton dynamics and parameterizing food-web models, yet their variation during viral infection remains poorly quantified. Here, we provide the first quantitative estimates of carbon and nitrogen fluxes in diatom virocells, showing that infection does not simply reduce nutrient assimilation but differentially reshapes carbon and nitrogen fluxes. These findings highlight the value of combining multi-omics approaches with *in vivo* measurements of metabolic fluxes to resolve the ecological and biogeochemical consequences of viral infection. Given the central role of diatoms in marine carbon and nitrogen cycling, a key next step will be to determine how assimilated carbon and nitrogen are partitioned among host biomass, viral production, and release upon cell lysis. Extending this approach to natural *Mediolabrus* populations, diverse host–virus systems, and environmentally relevant conditions will be essential to capture the variability of virocell metabolism at the ecosystem scale. Ultimately, incorporating virocell-specific traits into biogeochemical models should help constrain how viral infection alters marine primary production, nutrient cycling, and the efficiency of the biological carbon pump (47).

## Acknowledgements

The authors gratefully acknowledge the Roscoff Culture Collection (RCC) for providing the strains used in this study. We also thank the core facilities for mass spectrometry (METABOMER), flow cytometry (RECYF), and microscopy (Merimage) at the Station Biologique de Roscoff (EMBR-C, Sorbonne Université) for their support and assistance in method optimization. This work was funded by ANR-22-CE01-0018 BONUS (Biogeochemical consequences of diatom infection by marine viruses) and the Institut de l’Océan. Finally, we are grateful to the members of the BONUS consortium, particularly Marine Landa, for stimulating discussions and valuable insights.

