## Supplemental Information for "RNA virus infection reshapes carbon and nitrogen partitioning in a marine diatom"

**Supplementary information**

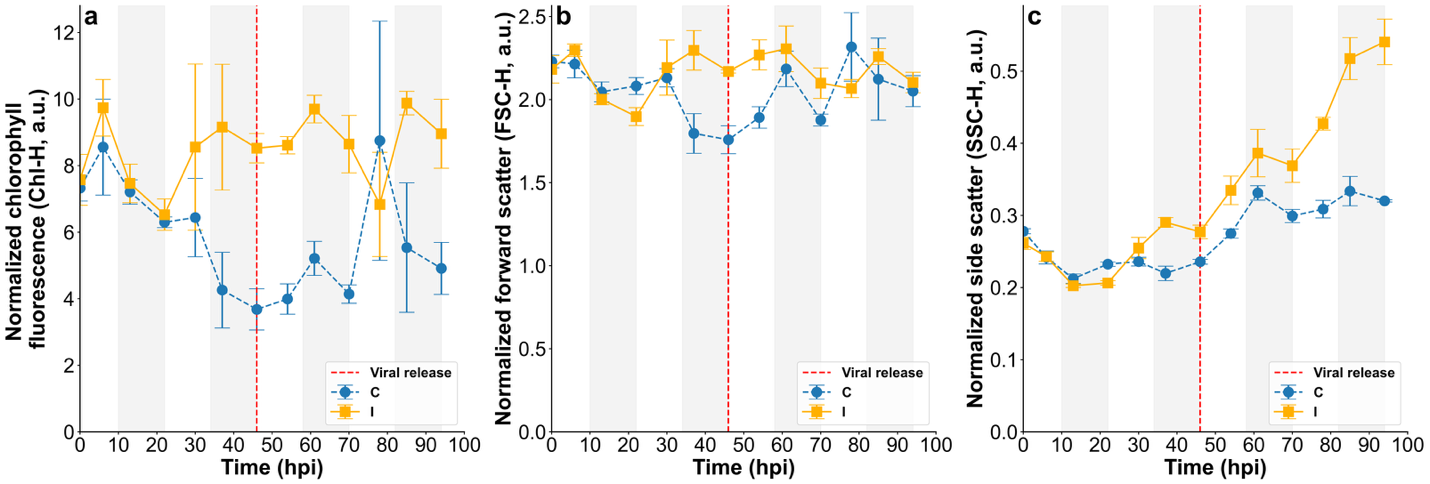

**Figure S1.** Variation in cellular optical properties **(a)** Normalized chlorophyll fluorescence (Chl-H), proxy of cellular chlorophyll content **(b)** Normalized forward scatter (FSC-H), proxy of cell biovolume (**c)** Normalized side scatter (SSC-H), proxy of intracellular structural complexity of *Mediolabrus comicus* over the course of the experiment. Control cultures (C) are shown by dashed blue line and Infected (I) cultures by solid yellow line. Values represent mean ± SD (n = 3). Grey boxes indicate night periods. The vertical dashed red line indicates the estimated timing of virus release, determined by plaque assay (FigS4.b).

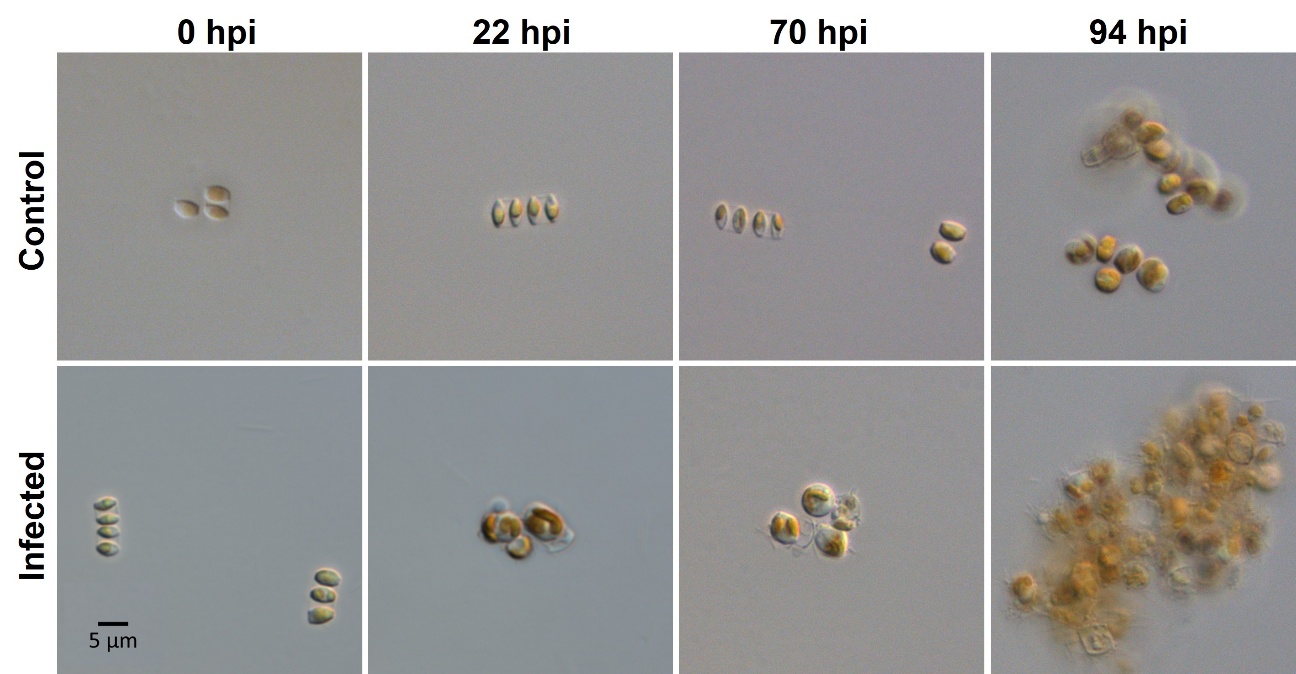

**Figure S2. Optical microscopy monitoring of Mediolabrus comicus** in control (C, top row) and infected (I, bottom row) cultures at 0, 22, 70, and 94 h post-infection. Infected cells tended to round up from 22 h post-infection onward and formed larger aggregates than those observed in the control cultures.

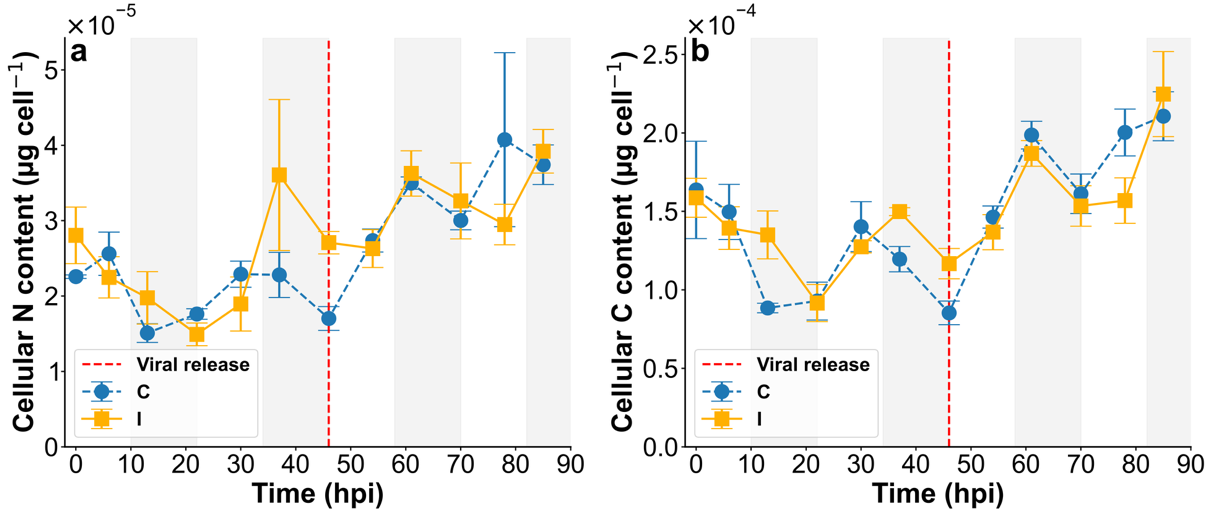
**Figure S3.** Variation of cellular content of **(a)** N (µg cell-1) and **(b)** C (µg cell-1) of *Mediolabrus comicus* over the course of the experiment. Control cultures (C) are shown by dashed blue line and Infected (I) cultures by solid yellow line. Values represent mean ± SD (n = 3). Grey boxes indicate night periods. The vertical dashed red line indicates the estimated timing of virus release, determined by plaque assay (FigS4.b).

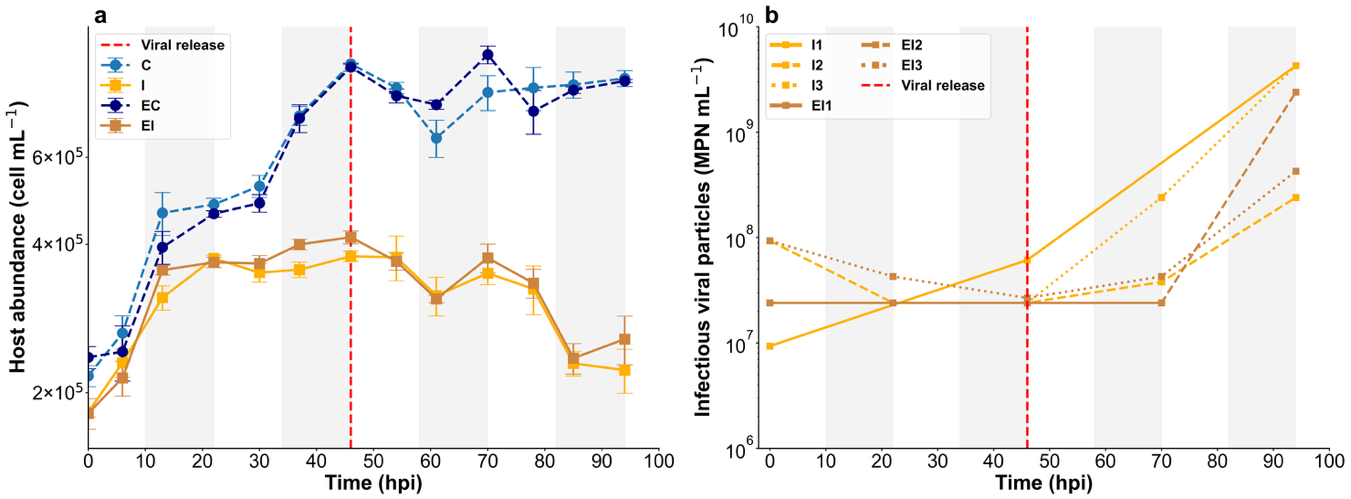

**Figure S4.** Effect of labeled isotope enrichment on infection dynamics of *Mediolabrus comicus*. **(a)** Variation of *M. comicus* cell abundance (cell mL^-1^) and **(b)** viral titer (infectious viruses mL-1). Control cultures (C) are shown by dashed blue line, Infected (I) cultures by solid yellow line in the experimental setup with non-isotopic enriched medium and Enriched Control (EC) are shown in dashed dark blue line, Enriched Infected (EI) in solid brown line amended with ^13^C-enriched NaH¹³CO₃ and ^15^N-enriched ¹⁵NH₄Cl assessed by isotope ration mass spectrometry. Values represent mean ± SD (n = 3). Grey boxes indicate night periods. The vertical dashed red line indicates the estimated timing of virus release, determined by plaque assay (FigS4.b).

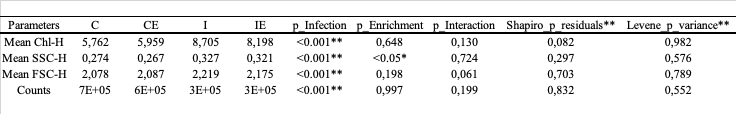

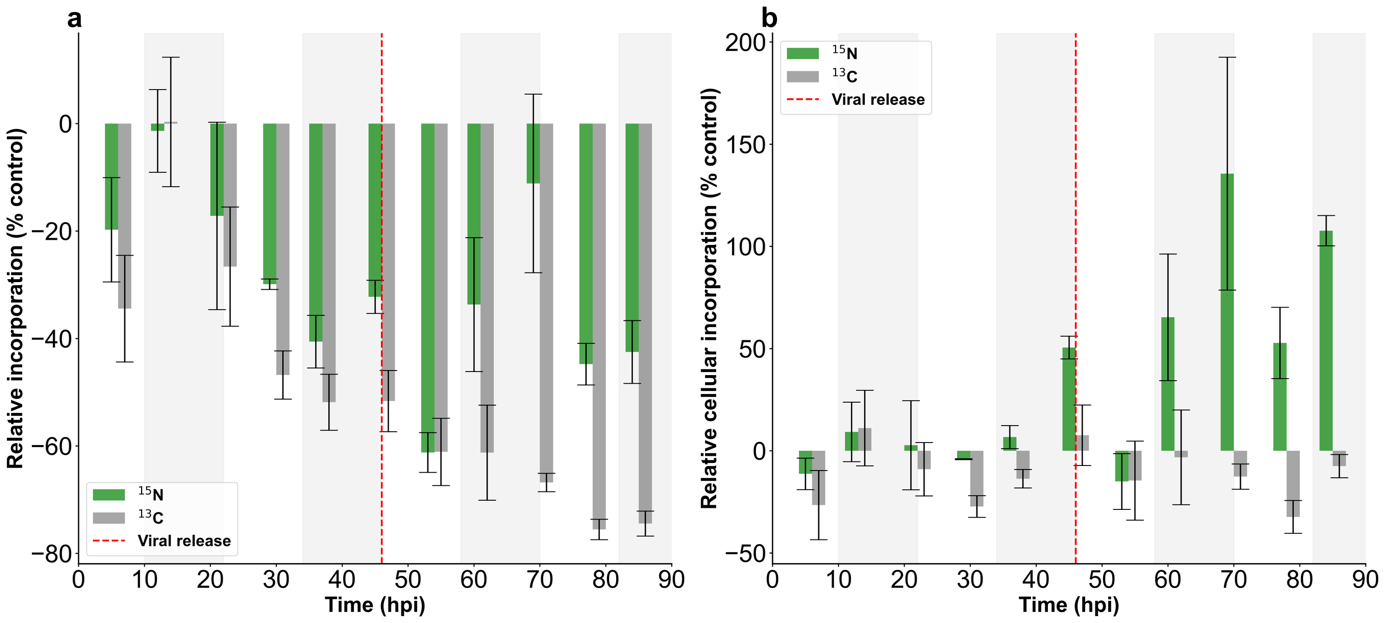

**Table S1.** Mean values of flow cytometry parameters: Chlorophyll content (Chl-H), Side Scatter (SSC-H, proxy for granulometry), and Forward Scatter (FSC-H, proxy for cell size), normalized to bead values; cell counts are expressed in cells·mL⁻¹. Data are shown for the four experimental conditions (Control Non-Enriched, Control Enriched, Infected Non-Enriched, Infected Enriched). Two-way ANOVA was performed at each timepoint on log-transformed data to test the effects of Infection, Enrichment, and their interaction. Reported p-values indicate significant effects of Infection (**p < 0.01**) and a slight effect of Enrichment on SSC-H (p 0.04). Assumptions of ANOVA were verified on residuals: normality by Shapiro–Wilk and homogeneity of variance by Levene’s test (p > 0.05 for both).

**Figure S5.** (a) Relative carbon (grey) and nitrogen (green) incorporation at the population-scale in infected cultures compared with control cultures, expressed as the percentage difference (%) over time and cumulatively from time 0. (b) Relative carbon and nitrogen incorporation at the cell level, expressed as the percentage difference (%) between infected and control cultures over time. The red dashed line indicates the onset of viral release. Values represent mean ± SD (n = 3). Grey boxes indicate night periods.
